# An amino-terminal fragment of apolipoprotein E4 leads to behavioral deficits, increased PHF-1 immunoreactivity, and mortality in zebrafish

**DOI:** 10.1101/2022.07.07.499128

**Authors:** Madyson M. McCarthy, Makenna J. Hardy, Saylor E. Leising, Alex LaFollette, Erica S. Stewart, Amelia S. Cogan, Tanya Sanghal, Katie Matteo, Julia T.T. Oxford, Troy T. Rohn

**Author notes:** **Correspondence:** Troy T. Rohn.

## Abstract

Although the increased risk of developing sporadic Alzheimer’s disease (AD) associated with the inheritance of the apolipoprotein E4 (*APOE4*) allele is well characterized, the molecular underpinnings of how ApoE4 imparts risk remains unknown. Enhanced proteolysis of the ApoE4 protein with a toxic-gain of function has been suggested and a 17 kDa amino-terminal ApoE4 fragment (nApoE4_1-151_) has been identified in post-mortem human AD frontal cortex sections. Recently, we demonstrated *in vitro*, exogenous treatment of nApoE4_1-151_ in BV2 microglial cells leads to uptake, trafficking to the nucleus and increased expression of genes associated with cell toxicity and inflammation. In the present study, we extend these findings to zebrafish (*Danio rerio*), which is an emerging *in vivo* model system to study AD. Exogenous treatment of nApoE4_1-151_ to 24-hour post-fertilization for 24 hours resulted in significant mortality. In addition, developmental abnormalities were observed following treatment with nApoE4_1-151_ including improper folding of the hindbrain, delay in ear development, deformed yolk sac, enlarged cardiac cavity, and significantly lower heart rates. Decreased presence of pigmentation was noted for nApoE4_1-151_ treated fish compared with controls. Behaviorally, touch-evoked responses to stimulus were negatively impacted by treatment with nApoE4_1-151_. A similar nApoE3_1-151_ fragment that differs by a single amino acid change (C>R) at position 112 had no effects on these parameters under identical treatment conditions. Additionally, triple-labeling confocal microscopy not only confirmed the nuclear localization of the nApoE4_1-151_ fragment within neuronal populations following exogenous treatment, but also identified the presence of tau pathology, one of the hallmark features of AD. Collectively, these *in vivo* data demonstrating toxicity as well as sublethal effects on organ and tissue development support a novel pathophysiological function of this AD associated-risk factor.

## 1. Introduction

Alzheimer’s disease (AD) is a neurodegenerative disease encompassing the most prevalent form of dementia characterized by amyloid plaques and neurofibrillary tangles (NFTs) [1, 2]. Early-onset AD has been associated with autosomal-dominant mutations in the amyloid precursor gene (APP), presenlin-1 and -2 (PSEN1/PSEN2) genes [2]. These mutations collectively comprising what is known as early-onset AD, affect approximately 5% of all known AD cases [3]. The majority of AD cases are characterized as late-onset in which the greatest risk factors for the disease are environmental (*e*.*g*., aging and lifestyle choices) in addition to the inheritance of an apolipoprotein (*APOE*) allele, namely apolipoprotein E4 (*APOE4*) [2]. The *APOE* gene has several isoforms of importance that are affected by a cysteine to arginine polymorphism: *APOE2* (C112, C158), *APOE3* (C112, R158), and *APOE4* (R112, R158) [4, 5]. A carrier of the *APOE4* allele increases the risk of developing AD by four-to twelve-fold [2]. However, the mechanisms of how ApoE4 contributes to increased risk of disease have remained elusive.

The *APOE* gene encodes the main cholesterol transporter protein (ApoE) in the CNS that is that is taken up by cells primarily through the low-density lipoprotein receptor (LDLR) family [6, 7]. Cholesterol transport in both the periphery and the CNS are vital for basal cellular function, but neurons are in critical need of adequate supply for synaptogenesis and neurite outgrowth [8]. The ApoE isoforms differ in their functional ability through the stepwise change in cysteine (C) to arginine (R) from ApoE2 to ApoE4 as described above [5]. The C112→R112 mutation has the ability to alter the side chain orientation of ApoE4 compared to ApoE2 and ApoE3 through the formation of a salt bridge combining R112 to E109 [5, 8, 9]. Data supports changes in the conformational structure of the isoforms from the C→R substitutions creates an increased likelihood of the generation of toxic fragmentation [10]. The role of ApoE4 proteolysis as a possible mechanism underlying disease risk has been supported by the findings of 17-20 kDa ApoE4 fragments in the prefrontal cortex from post-mortem AD patient tissue that localize within NFTs [10-14].

We recently examined the role of an amino-terminal fragment of ApoE4 previously identified in the human AD brain utilizing cultured BV2 microglia cells. Our findings demonstrated that exogenous application of this amino-terminal fragment of ApoE4 (nApoE4_1-151_) in microglial cells resulted in uptake of nApoE4_1-151_, trafficking to microglial nuclei, and the expression of numerous genes associated with inflammation [15-17]. To expand on this work, the current study employed an *in vivo* zebrafish system to study the fragment in a complex organism. The zebrafish model system has increasingly been used to study neurobiological questions in vertebrates. Some of the many benefits to this model are the rapid growth cycle, high fecundity rates, transparency of embryogenesis via externally fertilized embryos, and the early generation of stereotyped visualizable behavior at embryonic stages [18-20].

In the present study, exogenous treatment of zebrafish embryos with nApoE4_1-151_ led to an increase in toxicity and other morphological abnormalities. In addition, there was a trend towards decreased motor activation in the nApoE4_1-151_ treated-embryos as well as enhanced PHF-1 immunoreactivity. The findings of this study suggest that the single amino acid polymorphism from ApoE3 to ApoE4 includes a toxic gain-of-function providing a possible link between harboring the *APOE4* gene and enhance risk associated with AD.

## 2. Materials and Methods

### 2.1 Synthesis of nApoE_1-151_ fragments

Generation of nApoE4_1-151_ and nApoE3_1-151_ including synthesis of plasmid, expression in *E. coli*, and purification (>85% purity) was contracted out to GenScript Inc. (Piscataway, NJ). An anti-6X His-tag at the C-terminal end was added to facilitate purification. The verification of proteins was confirmed by DNA sequencing, SDS PAGE, and Western blot by using standard protocols for molecular weight and purity measurements. The concentration of recombinant proteins was determined by Bradford protein assay with BSA as a standard. Characterization of nApoE4_1-151_ in our laboratory was assessed by ELISA and Western blot analysis utilizing an anti-His antibody as previously described [11]. A similar fragment to apoE3 (nApoE3_1-151_) was utilized in order to directly compare the differences in toxicity and nuclear localization between nApoE4_1-151_ and nApoE3_1-151_.

### 2.2 Zebrafish Embryo Care and Maintenance

Animal husbandry and colony maintenance is handled by the Boise State University vivarium system. All animal protocols are coordinated with the Boise State University IACUC committee in accordance with recommendations from the Zebrafish Information Network (Zfin), IACUC protocol #AC18-011 and #AC21-009. Embryos were reared at 28.5°C in an enclosed incubation unit until treatment with exogenous protein fragment.

### 2.3 Treatment of Embryos with exogenous nApoE_1-151_ fragments

Treatment of embryos began at 24 hpf during the 19-somite-prim-6 stage in accordance with the Kimmel Staging Series [20]. Zebrafish embryos that were staged at the time of treatment to be above 25 hours (prim-6) or below 19 hpf (20-somite stage) were excluded from experimentation. Incubation of embryos was accomplished using a mixture of E3 media (Cold Spring Harbor Protocols, recipe for E3 medium for zebrafish embryos; doi:10.1101/pdb.rec066449) with various concentrations of nApoE4_1-151_ or nApoE3_1-151_ protein fragments. Following staging assessments, embryos were allocated to an incubation chamber in an even distribution depending on experiment. Embryos were kept with a density of no more than 5 embryos in a minimum of 100 µl of E3 media + protein fragment per well in 96 well plates. Control groups were raised in identical conditions with the only difference being the absence of nApoE_1-151_ fragments present in E3 media. Embryos were then placed in an incubator for 24 hours of undisturbed incubation at 28.5°C before being collected for experimentation.

### 2.4 Live Imaging (Light Microscope)

Live observations were recorded on an EVOS M5000 Light Microscope using brightfield settings. Embryos were imaged at 4X for a general view of the embryo(s) and 10X or greater were used for tissue specific image acquisition. Mortality was assessed as described above by comparing the number of nominal labels (1= Alive, 0= Dead/Non-viable) given to each group. A tally was taken via a contingency table in R to provide a raw count for each individual treatment group. Normality and equal variance were tested for prior to analysis. Each group count was then tested against each other via a Chi-square analysis and displayed via a mosaic display to indicate statistical significance by shading and examination of the Pearson residual values.

#### 2.4.1 Morphological Assessments

A standardized scale was generated across multiple standards resulting in a developmental abnormality score (**Figure 1**). Embryos were scanned throughout the z-axis to identify internal flaws, verify pigmentation changes, and gauge organogenesis at specific stages in coordination with the Kimmel staging series as well as comparison to the non-treated controls.

**Figure 1.**
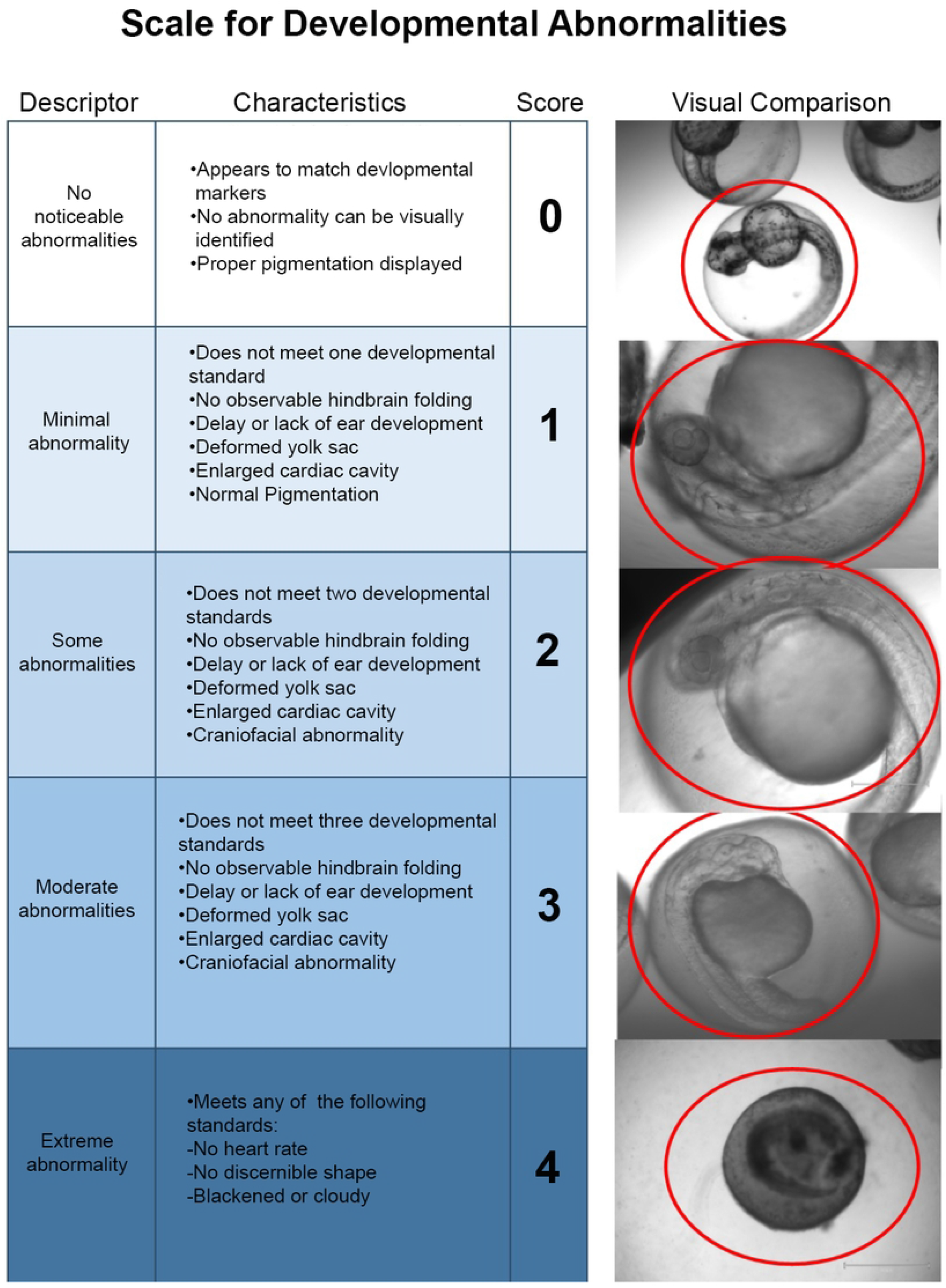
Semi-quantitative scale developed to assess morphological changes following treatment of zebrafish embryos with an amino-terminal fragment of nApoE4_1-151_. Rubric for developmental abnormalities was accomplished in 48 hpf zebrafish following 24-hour treatment with 25 µg/ml of nApoE41-151. This scale was established to quantify the effects of nApoE4_1-151_ as compared to untreated, control embryos. Identified hallmark defects that appeared consistently following treatment with nApoE4_1-151_ included inflation of pericardial cavity, enlarged hearts, pigmentation alterations, and delays or lack of development in ear and brain structures. Data are representative of 10 embryos treated with 25 µg/ml nApoE4_1-151_ per trial for a total of 30 embryos.

#### 2.4.2 Heart Rate Determination

Live imaging was also applied for mortality assessments of embryos in the manner of identifying heartrate over the course of 10 second intervals. If no heart rate was detected, stimulation of embryo through water movement was performed to verify lack of response to stimuli. If no response to stimulation was found in combination with a lack of heartbeat, the embryo was deemed non-viable. Heart rate was measured in a continuous manner by beats per minute (bpm) as described above. Samples were averaged by replicate to stabilize variation within groups. Five sets of heart rate collection dates were tested through two-way ANOVA modelling with three treatment groups (Control, nApoE3_1-151_, and nApoE4_1-151_) in R statistical software. ANOVA model was assessed for assumptions after creation of model in R.

### 2.5 Behavioral Assays

#### 2.5.1 Tail Flick Behavioral Test

Larvae at 72 hpf were acclimated to the testing area for 2 minutes prior to experimentation. Five-minute videos were taken of each larvae using a Motic MGT 101 Moticam recording device with an LED-60T-B light ring. Videos were recorded down each treatment group column, then across well rows in a 96-well petri dish. Immediately following testing, the samples were euthanized using IACUC and University Guidelines, preserved in 4% PFA/PBT, then stored in 100% ethanol at -20°C. During each recording, individual larva was documented using a numerical code to designate treatment groups for a single-blind procedure, in which one researcher recorded and encoded video names for treatment and a second researcher analyzed coded videos. Scoring was completed by examining videos while documenting the number of spontaneous tail flicks in total for each larva over a 5-minute span. A single, spontaneous tail flick was determined by counting the number of times each larva bent their tail away from center and returned to the center axis. Samples were prepared by averaging 5 tail flicks per replicate over the course of 5 replicates with both nApoE3_1-151_ and nApoE4_1-151_ at a final concentration of 10 µg/ml. Ten µg/ml was used as a concentration that would evoke behavioral responses without a decrease in survivability or any visible morphological deficits. Data analyzed for each trial represent the averaged results for each collection date. Data was analyzed through two-way ANOVA modeling with three treatment groups (Control, nApoE3_1-151_, and nApoE4_1-151_) in R statistical software. The ANOVA model was assessed for assumptions after creation of model in R.

#### 2.5.2 Touch Evoked Movement Response Assay (TEMR)

Individual larvae at 72 hpf were moved to a 14×14 mm round glass bottom petri dish filled with E3 Media 2 minutes prior to testing for acclimation under light conditions for testing. Two-minute videos were recorded on Motic MGT 101 Moticam recording device with an LED-60T-B light ring. After the start of the recording at time (T=0), embryos were tapped lightly with a blunt probe every 15 seconds. Directly following testing, larvae were euthanized using IACUC and University Guidelines, preserved in 4% PFA/PBT, then stored in 100% ethanol at -20°C. Videos were analyzed in Noldus Behavioral Software version 15.0. Criteria for scoring was based on the number of responses following the evoked stimulus. Data was grouped by treatment and analyzed in R statistical software. Data was assessed for normality and variance. The number of responses to the evoked stimulus was analyzed using One-Way ANOVA (aov; car package). A non-parametric Kruskal-Wallis test was applied to duration data as the data was not normal.

### 2.6 Immunofluorescence labeling

For all immunohistochemical procedures in this study, a standard protocol was followed for tissue preparation, and preparation of the slides. At the conclusion of treatment experiments, embryos were sacrificed prior to manual removal of chorion. Following de-chorionation, embryos were fixed in 4% paraformaldehyde (PFA)/phosphate buffered saline (PBS) overnight at 4°C and prepared for 5 µm paraffin-embedded sectioning using a Leica RM2235 Microtome at 4 °C. Paraffin-embedded sections were used for staining following rehydration and a series of washes in PBS with tween-20 0.05% (PBST). Blocking of sections for non-specific staining was accomplished using a standard incubation buffer consisting of 1% normal goat serum, 2% bovine serum albumin in PBST for 2 hours at room temperature. Primary antibodies, detailed in **Table 1**, were incubated with slides for 18-24 hours at 4°C. Sections were then washed in triplicate for 5 minutes with PBST. Slides were then incubated in the appropriate secondary antibody for 1 hour at room temperature. Following the final wash, DAPI infused soft mount was placed on slides and allowed to set before coverslip addition. Sections were analyzed for preliminary staining with EVOS M5000 light cubes at lowest intensity settings. Sections were then kept in the dark at 4°C until used for confocal imaging. Following labeling, confocal assessment of the localization of nApoE4_1-151_ in neuronal cell populations was as previously described [17]. All images and z-stacks generated were obtained using Zeiss Microscope, LSM 510 Meta confocal imaging system (Carl Zeiss, Oberkochen, Germany) and processed using Zen blue edition (Carl Zeiss, Göttingen, Germany).

**Table 1.**
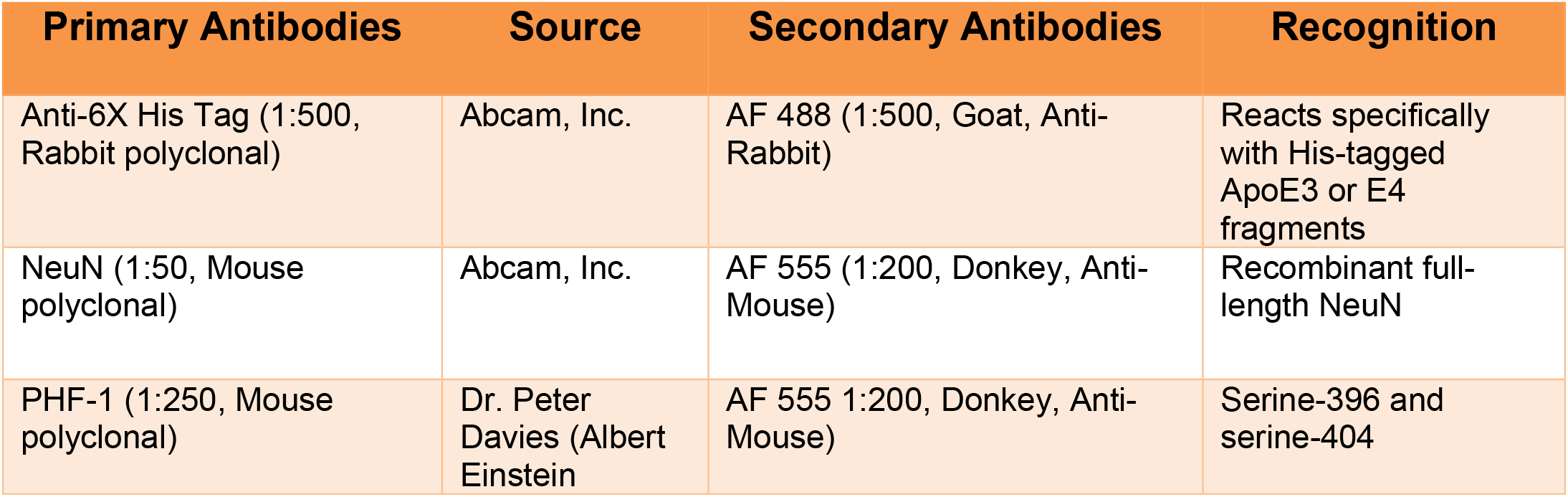
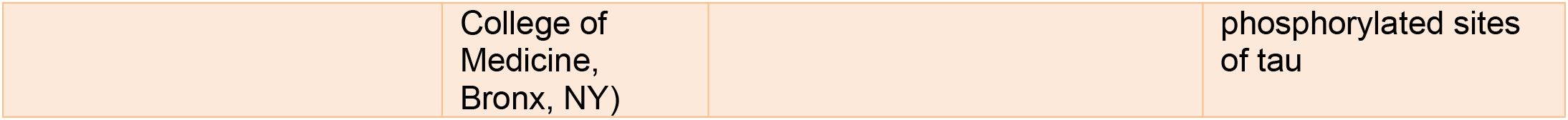
Description of antibodies used for immunofluorescence experiments.

## 3. Results

We chose to examine the role this specific amino-terminal fragment of ApoE41-151 for several reasons. First, we documented widespread evidence for this fragment in the human AD brain, where it localized within nuclei of microglia [17]. Second, generation of this fragment was documented following incubation of full-length ApoE4 with matrix metalloproteinase-9 (MMP-9) [17]. Finally, Recent data from our lab suggests that, *in vitro*, a 151 amino-terminal fragment of ApoE4 (nApoE4_1-151_) can traffic to the nucleus leading to toxicity and expression of inflammatory genes in BV2 microglia cells [15, 16, 21]. The purpose of this current study is to expand those findings *in vivo*, by assessing the mechanisms by which this fragment may induce toxicity, developmental abnormalities, and behavior deficits in a model system consisting of zebrafish.

### 3.1 Morphological Assessments

A semi-quantitative morphological assessment revealed treatment group specific phenotypes that were predominantly present in the nApoE4_1-151_-treatment group. A scale was generated across multiple standards resulting in a developmental abnormality score (Fig. 1).

Embryos were scanned throughout the z-axis to identify internal flaws, verify pigmentation changes, and gauge organogenesis at specific stages in coordination with the Kimmel staging series as well as comparison to the non-treated controls. Control groups were used to set a standard of comparison of which nApoE3_1-151_ followed closely in most regards except for loss of pigmentation in both treated groups (blue arrows, Fig. 2A-C). Features most commonly observed for nApoE4_1-151_ groups in addition to loss of pigmentation were lack of hindbrain folding at cerebellar primordium (orange arrows, Fig. 2A-C), as well as enlargement of the cardiac cavity (red arrow, Fig. 2C). Quantification of morphological abnormalities are depicted in Fig. 2D based on our standardized scale, with nApoE4_1-151_ showing the most severe degree of changes following treatment with a significant increase in developmental abnormality scores compared to untreated controls. Heart rate measurements reported a significant

**Figure 2.**
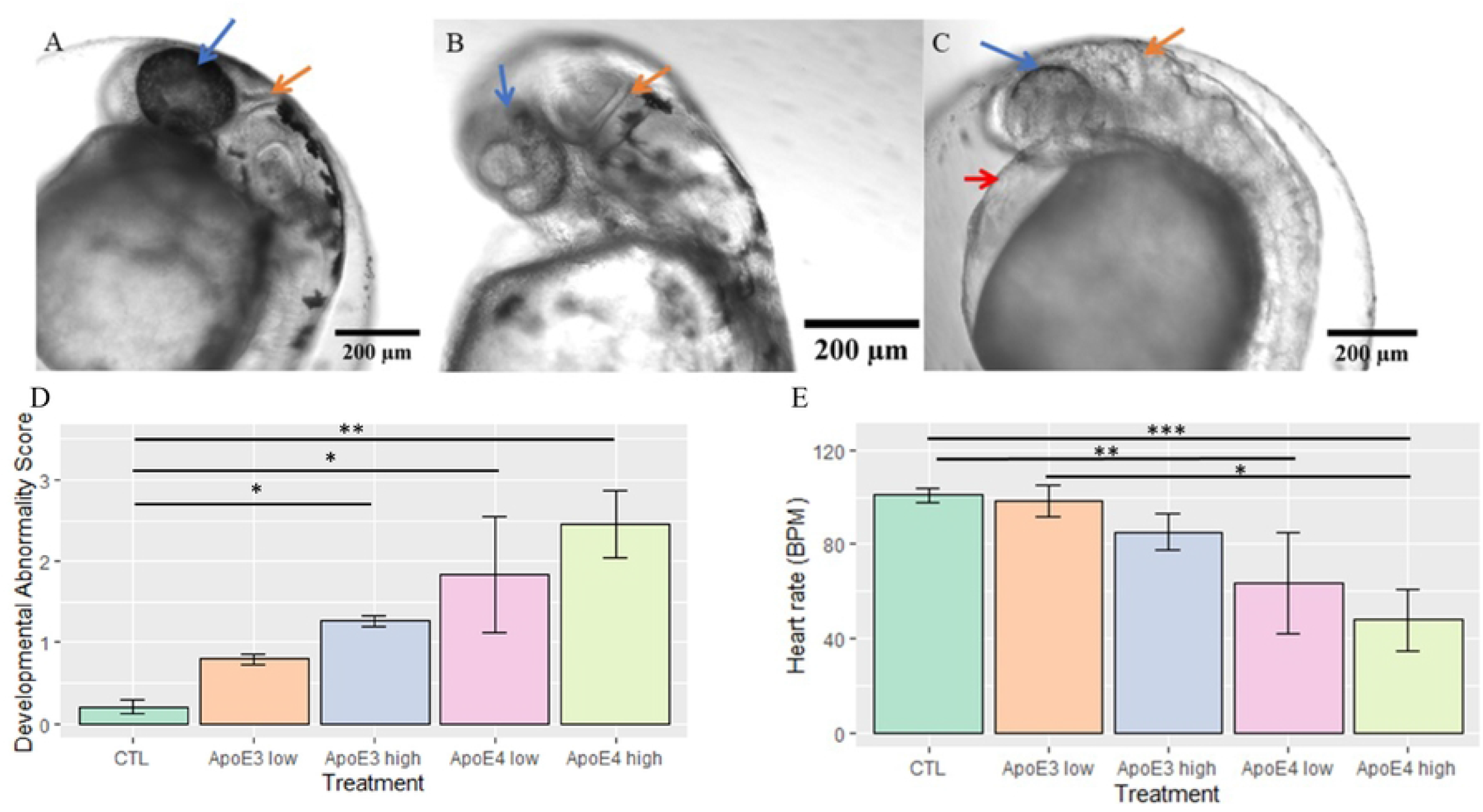
A sublethal concentration of nApoE4_1-151_ leads to morphological abnormalities in zebrafish embryos at the hatching phase. Representative light phase contrast microscopic images following live imaging of embryos at 48 hpf following a 24-hour period with respective treatments (Control, 25 µg/ml of nApoE3_1-151_, or 25 µg/ml of nApoE4_1-151_. **A:** Arrows point to consistent morphological changes as compared to untreated controls that are healthy and categorized as having a developmental abnormality score of <1.0 (Panel A). The blue arrows designate pigmentation changes; orange arrows designate cerebellar primordium junction differences in the hindbrain. **B:** Exogenous treatment with nApoE3_1-151_ impacted pigmentation pattern in otherwise healthy embryos. Embryos in this category were most likely to receive a score of <1. **C:** 24-hour incubation of a sublethal concentration of nApoE4_1-151_ resulted in developmental abnormality scores >3. Embryos in this category were typically observed to be delayed in development with limited hindbrain folding (orange arrow), limited or lacking pigmentation (blue arrow), as well as enlargement of the cardiac cavity (red arrow). **D**. Quantitative developmental abnormality scores for each treatment group following treatment of embryos for 24 hours with respective fragments at low E3 (orange bar), E4 (pink bar) concentrations (25 µg/ml) or at high concentrations (50 µg/ml) E3 (gray bar), E4 (yellow bar). The nApoE4_1-151_ 25 µg/ml-treated groups were significantly different from controls (H(4)=-2.43, *p*=0.0074). At 50 µg/ml both nApoE4_1-151_ (H(4)=-3.32, p=0.0004) and nApoE3_1-151_-treatment groups (H(4)=-1.77, p=0.037) were significantly different from controls. Errors bars represent ± S.E.M. *p<0.05, **p<0.01, ***p<0.001 **E**. Heart rate data obtained from live microscope analyses in 25 µg/ml and 50 µg/ml treatment groups nApoE3_1-151_ and nApoE4_1-151_ compared to non-treated controls. nApoE4_1-151_ (pink bar, 25 µg/ml) was significantly different from controls (H(4)=1.77, *p*=0.038). nApoE4_1-151_ (yellow bar, 50 µg/ml) was significantly different from nApoE3_1-151_ 25 µg/ml (H(4)=1.94, p=0.026) and controls (H(4)=2.665, p=0.0036). Errors bars represent ±S.E.M. All other comparisons were insignificant. *p<0.05, **p<0.01, ***p<0.001

### 3.2 Survivability and Mortality Assessments

Survivability curves indicated that treatment of zebrafish embryos at 24 hpf lead to significant mortality. There was a 90-100% reduction in viable embryos 48 hours after treatment began (Fig. 3A). In addition, nApoE4_1-151_ treated-embryos failed to recover after removal of treatment media. The survivability of embryos following treatment with 25 µg/ml of nApoE4_1-151_ was reduced by 50% within 24 hours of treatment (yellow dotted line), whereas at 50 µg/ml, it was reduced by 50% by 12 hours (green dotted line, Fig. 3A). As a control, we also tested the impact of nApoE3_1-151_ on survival. In this case, identical concentrations of nApoE3_1-151_ had little impact on embryo survival following treatment even at the highest concentration of 50 µg/ml (<15%) (blue dashed line, Fig. 3A).

**Figure 3.**
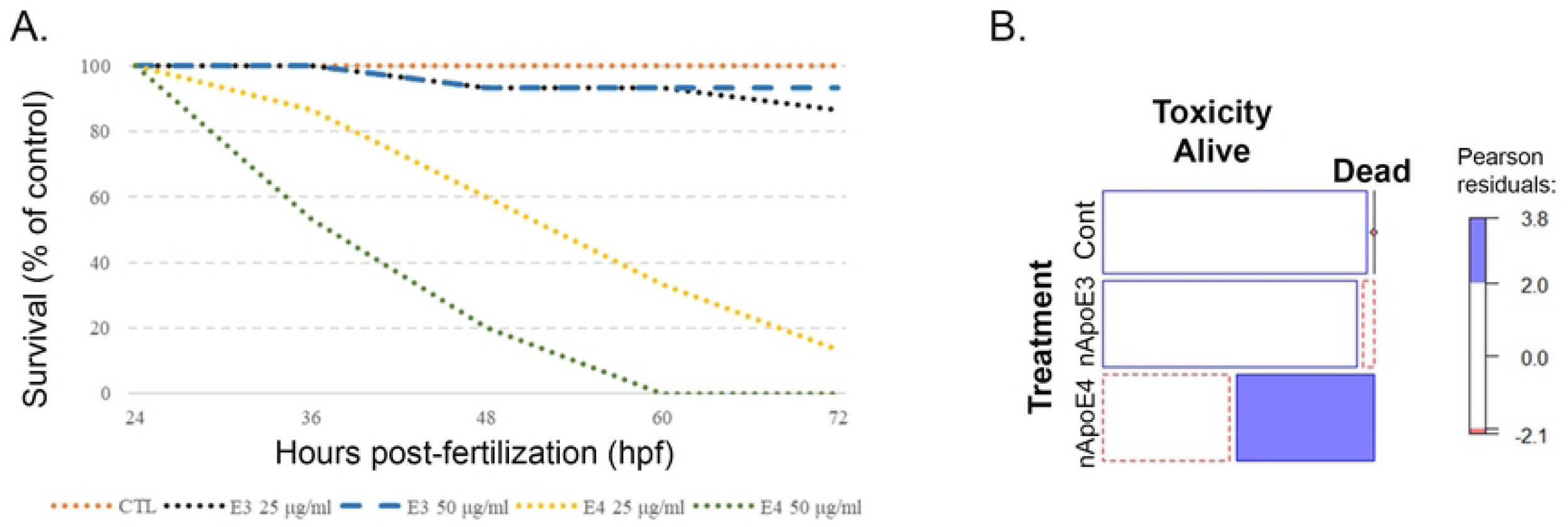
Survivability is decreased in zebrafish embryos following exogenous treatment with an amino-terminal fragment of nApoE4_1-151_. **A.** Embryos at 24 hpf (prim-9 stage) were segregated into three groups: controls (untreated), 25 μg/ml or 50 μg/ml nApoE3_1-151_, and 25 μg/ml or 50 μg/ml nApoE4_1-151_. Embryos that lacked a heartbeat for 10 seconds were stimulated to induce movement. If no movement or heartbeat was detected, embryos were considered to be non-viable. Significant mortality was observed at both concentrations of nApoE4_1-151_ by 48 hpf (orange and green dotted lines) compared to non-treated controls (orange dotted line) or nApoE31-151 (blue dashed line and black dotted lines). N=3 independent trials, N=5 fish/treatment. **B.** The Mosaic Plot depicts mortality based on a lack of heartbeat and response to physical stimuli following exogenous treatment of 48 hpf zebrafish embryos with either 25 µg/ml nApoE3_1-151_ or nApoE4_1-151_ for 24 hours. The blue filled region of the bar graph designates 25 µg/ml treatment of nApoE4_1-151_ which led to a significant portion of the embryos being designated as dead. The red dotted portion of the bar graphs indicates less than expected were alive (p=1.17e-13). All blank cells indicate the sample group followed the estimated trend. Data indicated significant morality for only the nApoE4_1-151_ group. N=3 independent experiments, 15 embryos/treatment.

Mortality data reported a high nApoE4_1-151_ associated mortality compared to both nApoE3_1-151_ and non-treatment groups (p-value compared to non-treated controls was 1.17e^-13^) (Fig. 3B). nApoE3_1-151_ and controls revealed a nearly identical mortality rate with <10% drop in mortality for nApoE3_1-151_ compared to untreated controls. These results suggest that changing a single amino acid (C>R) at position 112 is sufficient to induce significant toxicity when zebrafish embryos are treated exogenously with nApoE4_1-151_.

### 3.3 Neuronal localization of nApoE4_1-151_ and the presence of tau pathology

To assess the cellular localization of nApoE4_1-151_ following treatment of zebrafish embryos, confocal analysis was undertaken following fixation and sectioning of treated embryos. To track potential nApoE4_1-151_ uptake, we utilized an anti-6X His-tag antibody (1:500), together with DAPI (nuclear stain) and markers for neuronal cells including Neuronal Nuclei (NeuN). NeuN is part of the RNA splicing machinery and is predominantly found in the nucleus of post-mitotic neurons [22]. As a comparison, we also performed identical studies tracking the localization of nApoE3_1-151_. As depicted in Figure 4, nApoE4_1-151_ is taken up by neurons following exogenous treatment of 48 hpf zebrafish embryos that colocalized with NeuN (arrows, Fig. 4D). Of interest was the punctate staining of nApoE4_1-151_ which supports our previous staining pattern observed in transformed BV2 microglial cells, suggesting possible aggregation of the nApoE4_1-151_ fragment [17]. In contrast, although nApoE3_1-151_ was detected within neurons, the staining pattern was different: in this case, perinuclear staining of nApoE3_1-151_ was evident and there was very little co-localization with NeuN (arrows, Fig. 4H. These data also support our previous findings in that nApoE3_1-151_ namely is confined within the cytoplasm of cells with little trafficking to the nucleus [17]. The specificity of both fragments was confirmed following the results in non-treated controls that showed a complete lack of anti-His staining (Fig. 4C).

**Figure 4.**
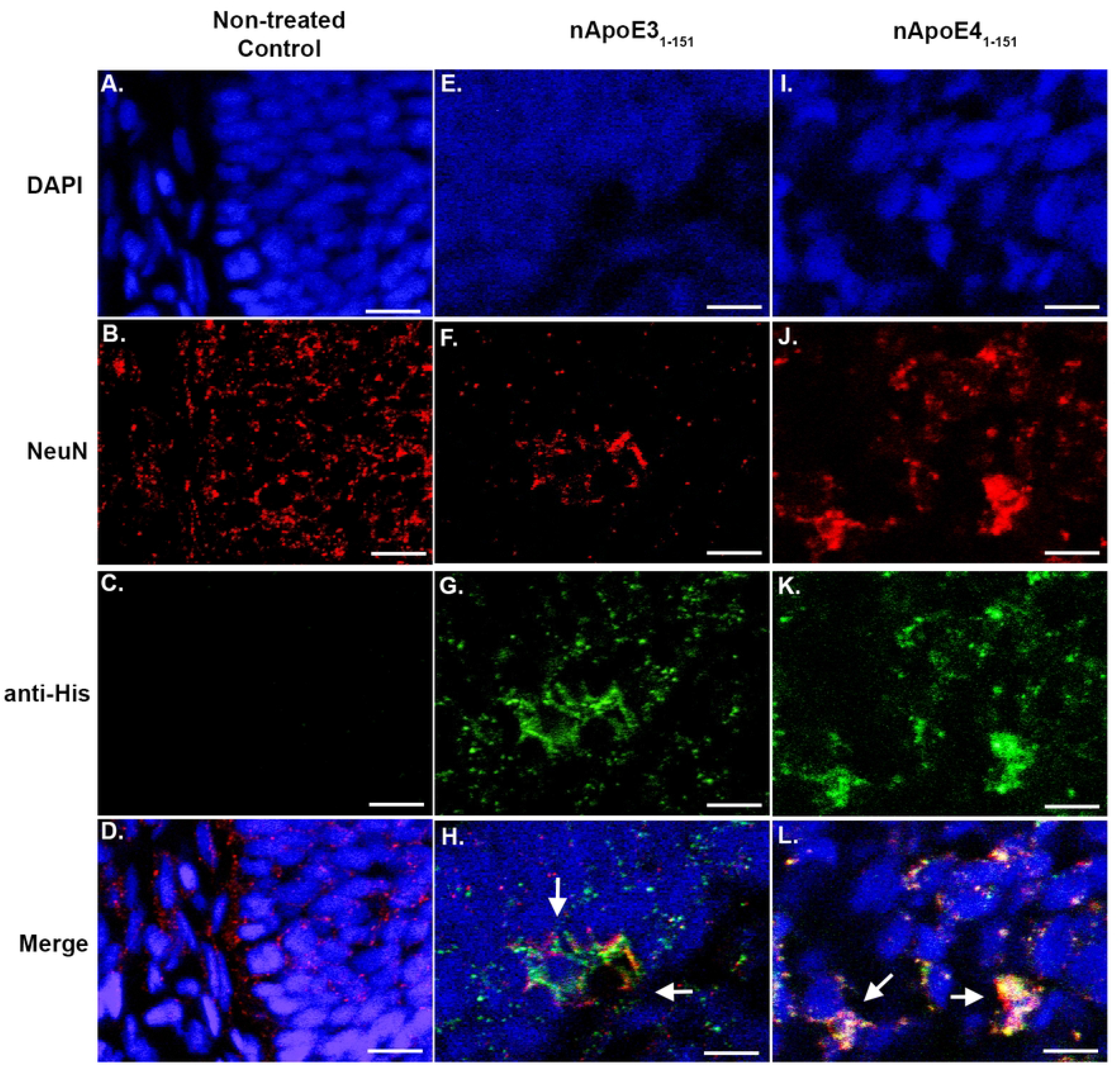
Exogenous treatment of zebrafish embryos with amino-terminal fragments of nApoE41-151 leads to nuclear localization. **A-D**. Representative images from confocal immunofluorescence in 5 mm paraffin-embedded sections of non-treated control 48 hpf zebrafish embryos that were stained with DAPI (A), NeuN (1:50) (B), anti-His antibody (1:500) (C), and the merged image in Panel (D). There was no detection of any nApoE?_1-151_ fragments in untreated control neuronal cells as indicated by the lack of labeling in Panel C. **E-H**. Identical to Panels A-D with the exception that embryos were exogenously treated for 24 hours with 25 µg/ml of nApoE3_1-151_. Perinuclear localization of nApoE3_1-151_ was evident (Panel G and H, arrows) under these conditions. **I-L**. Parallel experiments as in Panels E-H with the exception that zebrafish embryos were treated exogenously for 24 hours with 25 µg/ml of nApoE41-151. In this case, strong nuclear localization of nApoE4_1-151_ was evident within neuronal cells (Panels K and L, arrows). For both fragments, staining appears punctate and co-localized with NeuN and DAPI. All images were captured within the area of the cerebellum and fourth ventricle. Data are representative of three independent experiments. All scale bars represent 10 µm. 12

Additional immunofluorescence studies were undertaken to assess any potential relevance to known AD pathology. Previous studies have demonstrated that amino-terminal fragments of ApoE4 localize in NFTs of the human AD brain [11] and may induce neurofibrillary changes in cultured neurons [10]. Zebrafish are known to express gen orthologues to the human *MAPT* gene including *MAPTA* and *MAPTB* [23]. Therefore, we examined whether treatment of zebrafish embryos with nApoE4_1-151_ led to similar pathological changes to tau by confocal IF using PHF-1 that recognizes hyperphosphorylated, fibrillar forms of tau present in the human AD brain. At 48 hpf, we were able to detect PHF-immuno-reactivity within the brain of zebrafish embryos that was not present in the untreated controls or in nApoE3_1-151_-treated embryos (**Supplemental Figure 1**). Because PHF-1 staining was relatively weak at this time point, we expanded this experiment to include treated embryos at 72 hpf. Similar results were obtained, however, much more robust labeling of PHF-1 within apparent neurons was evident under these experimental conditions (arrows, Fig. 5I-L). In Figure 5K, a merged, high-magnification of another representative treated embryo is shown. In this case, PHF-1 labeling appeared fibrillar in nature, which is characteristic of PHF-1 staining within NFTs of the AD brain [24]. These results support the linkage of nApoE4_1-151_ to one of the significant hallmark pathologies found in AD, namely neurofibrillary tangles.

**Figure 5.**
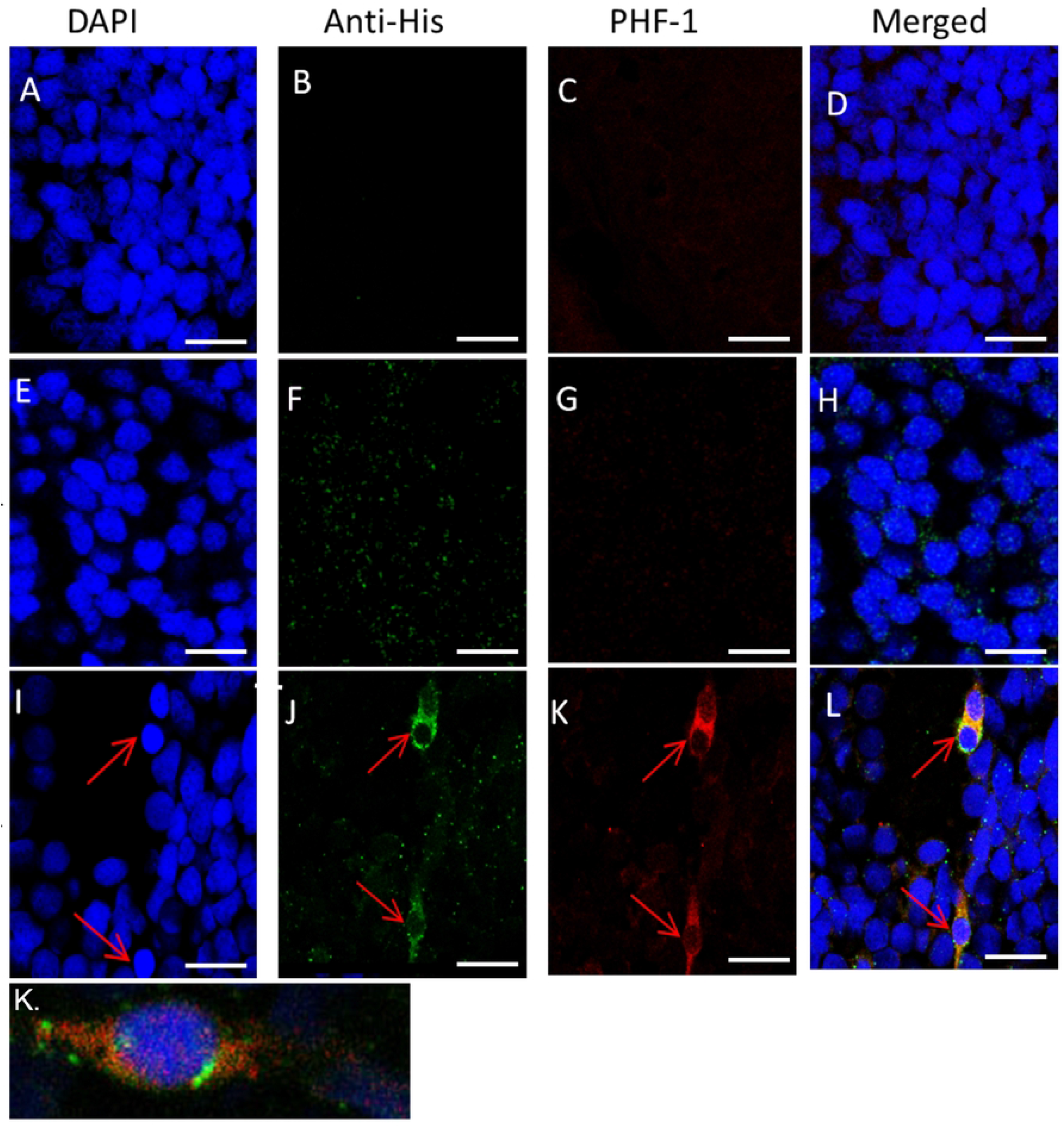
Tau pathology present after treatment with exogenous nApoE4_1-151_ fragment in 72 hpf zebrafish brain. Representative 40X images from confocal immunofluorescence in 5 mm paraffin embedded sections of non-treated control 72 hpf zebrafish that were stained with DAPI (A-I), Anti-6X His antibody to detect nApoE4_1-151_ (B-J), PHF-1 (C-K) and merged images for all three markers (D-L). As expected, there was little staining observed for of nApoE4_1-151_ or PHF-1 (A-D). Strong PHF-1 labeling was only observed following treatment with nApoE4_1-151_ (I-L). Panel K depicts a separate, representative merged image following treatment with nApoE4_1-151_. In this case, at high magnification the fibrillar nature of PHF-1 labeling was apparent.

### 3.4 Motor deficits in juvenile zebrafish following treatment with a sublethal concentration of nApoE4_151_

As an initial approach, we assessed whether a stereotypical motor behavior in zebrafish, spontaneous tail flicking, was diminished following treatment with nApoE4_151_ or nApoE3_151_ at a concentration that does not generate any mortality or visible morphological deficits (10 µg/ml). Figure 6A depicts the results of this experiment showing that zebrafish treated with nApoE4_1-151_ performed the least number of tail flicks per group with >50% reduction from untreated controls. Treatment of zebrafish larvae with nApoE3_1-151_ performed similarly to controls. However, no difference was significant across any group (*F* (2,27) = 1.24, *p*=0.305). It’s noteworthy that control zebrafish demonstrated minimal spontaneous tail flick behavior and there were large variations within groups. The lack of response in even our control cohorts indicates that there is limited motor movement in 72 hpf in an unstimulated environment. Therefore, a second motor behavior task was undertaken whereby zebrafish were stimulated to move using a blunt instrument. This type of behavior is known as the touch-evoked movement response (TEMR) test and results mirrored the findings from the spontaneous tail-flick experiment (Figure 6B). In this case, control and nApoE3_151_-treated groups responded to 90% of applied stimulations. In contrast, nApoE4_151_-treated fish responded to fewer than 50% of evoked stimulations (**Supplemental videos 1 and 2**). However, as in Figure 6A, due to high variations within groups, no statistical significance was observed (F(12)=1.482, p=0.266. Taken together, our results show strong trends for motor behavior impairments following treatment of zebrafish with sub-toxic concentrations of nApoE4_151_, that were not present in non-treated controls nor in nApoE3_151_-treated fish.

**Figure 6.**
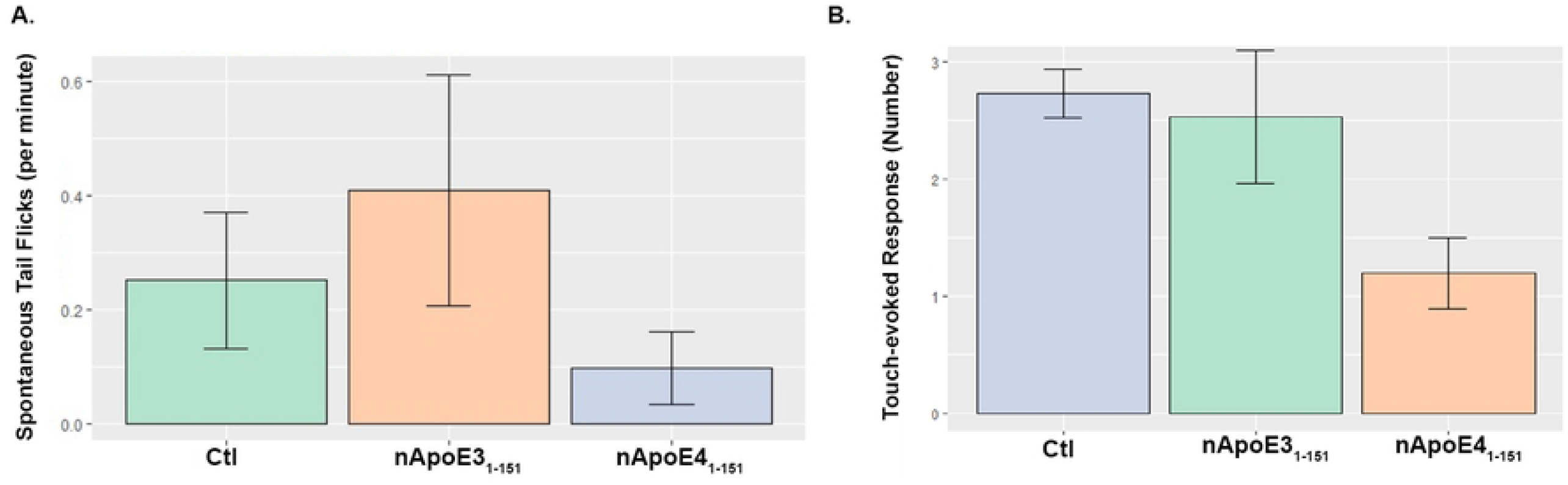
Negative trends in motor behavior in zebrafish following treatment with nApoE4_1-151_. **A**. Groups for non-treated controls (green bar), nApoE3_1-151_ 10 µg/ml (orange bar), and ApoE4_1-151_ 25 µg/ml (blue bar)) were assessed via video monitoring to determine number of spontaneous tail flicks per minute that were then averaged per group for each trial. Data are representative of N=5 trials, for a total of 25 embryos per group. Data depicted show limited spontaneous tail flick activation from every group with no difference detectable between groups (*F(*2,27)=1.24, *p*=0.305). **B**. Results from the touch-evoked response motor behavior experiment similar to (A). Non-treated controls had a 90% response rate to the evoked, tactile stimulus, whereas for nApoE4_1-151_-treated groups responded to fewer than 50% of stimuli. No significant difference was observed (*F*(12)=1.482, p= 0.266).

## 4 Discussion

Despite intensive research efforts, the pathophysiological relationship between harboring the *APOE4* allele and the development of late-onset AD remains largely unknown. This question is further complicated by the fact that only the ApoE4 protein represents a significant risk factor even though it differs from ApoE3 by a single amino acid at position 112 and ApoE2 by two amino acids (positions 112 and 158) [5]. Both substitutions lead to the replacement of a cysteine residue with an arginine residue [5]. One possible hypothesis leading to increased dementia risk is the propensity of the ApoE4 isoform to be highly susceptible to proteolysis compared to E3 and E2 [25]. Prior research from our lab supports the hypothesis that ApoE4 fragmentation via the metalloproteinase-9 (MMP-9) may contribute to AD pathology and inflammation [17]. Additional studies have also shown neurotoxic and pro-inflammatory responses to the fragmentation of ApoE4 [26-29]. Recently we identified a 151 amino-terminal fragment of ApoE4 (nApoE4_1-151_) that localized within the nucleus of both neurons and microglia cells of the human AD brain [17] and *in vitro*, we demonstrated this fragment is taken up by BV2 microglia cells, traffics to the nucleus and leads to the upregulation of thousands of genes, many of which associated with microglia activation and inflammation [15, 21]. In the current studies, we expanded these findings utilizing an *in vivo* model system consisting of zebrafish.

One of the more distinct advantages of the zebrafish is the optical clarity of the embryos allowing for the investigation throughout the developmental process using non-invasive imaging techniques. There is also a high degree of conservation between zebrafish and human brain organization including both neuroanatomic [30, 31], neurochemical [32], and behavioral circuitry including learning [33], touch [34], and decision making [35]. Moreover, zebrafish have served as excellent models to study the pathophysiology underlying AD including a study that demonstrated intraventricular injection of Aβ_1-42_ in the embryonic brain leads to memory loss and cognitive deficits along with increased tau phosphorylation [36, 37]. Taken together, the zebrafish presents itself as a novel model system to examine the potential effects of nApoE4_1-151_.

As an initial approach, we treated zebrafish embryos at 24 hpf and examined any potential morphological changes 24 hours later (48 hpf). Compared to untreated controls, nApoE4_1-151_ exposure led to changes to the nervous system, the heart, and pigmentation development. The nervous system was visibly impacted by the lack of hindbrain folding that is typical in 48 hpf zebrafish along the cerebellar primordium. Additionally, nApoE4_1-151_ treatments at both a low (25 µg/ml) and high (50 µg/ml) concentrations produced significant differences from controls in terms of developmental abnormalities and reduced heart rates. The morphological results of enlargement of hearts following nApoE4_1-151_ treatment are intriguing based on the well-known link between inheritance of *APOE4* and an increased risk of cardiovascular disease [38]. Confocal imaging revealed the colocalization of nApoE4_1-151_ with NeuN within the hindbrain region around the medulla oblongata which regulates heart rate. These data support the hypothesis that nApoE4_1-151_ may destabilize the hindbrain networks during the incubation period leading to downstream cardiovascular deficits, including a reduced heart rate.

From early stages of development, zebrafish swimming behavior and response to external stimuli can be assessed. The responses to external stimuli can be detected at early larvae states (72 hpf) in which zebrafish show escape response swimming behavior, for example in response to touch directed to either the head or tail [39]. Similar to the cardiovascular system, the touch-evoked response is under control of the hindbrain [40, 41]. Specifically, the touch escape response and spontaneous tail flicks are both proposed to be regulated by the Mauthner cells located in the zebrafish hindbrain [42, 43]. Treatment of zebrafish larvae with nApoE4_1-151_ led to less than half of the stimulation attempts whereas non-treated controls and nApoE3_1-151_ responded to over 90% and 80% stimulation attempts, respectively. Increased detection of nApoE4_1-151_ compared to controls within the hindbrain region could be the rationale for the effects observed in both of these locomotor assays. Presently, it is not known how nApoE4_1-151_ leads to these motor deficits or at what stage of this behavior does nApoE4_1-151_ interfere: initiation or execution of the behavioral response? Initiation of the response would imply the sensation of the stimulus was never received through the rohon-beard cells or the dorsal root ganglia which have been shown to share the responsibility of tactile sensation during the 72 hpf period [44].

Interestingly, nApoE3_1-151_ did not induce significant changes to morphology, toxicity, or locomotor behavior. The only major effect of nApoE3_1-151_ exogenous treatment was a decrease in pigmentation. During embryogenesis, pigment cell precursors migrate from the neural crest to generate an embryonic/early larval pigment pattern by approximately day 4 consisting of yellow xanthophores disperse over the flank and melanophores and iridophores along the dorsal and ventral edges of the body and over the yolk, and a few melanophores in a lateral stripe along the horizontal myoseptum [45]. Our current results indicating a significant decrease in pigmentation by both E3 and E4 fragments suggests a common mechanism that disrupts the normal pathway of pigment patterning. One possible mechanism is the disruption of the bone-morphogenetic protein (BMP) signaling pathway that when activated allows for the proper dorsoventral patterning and induction of the neural crest [46]. Further research is necessary to uncover the mechanism of pigmentation disruption by ApoE fragments, but our results suggest that they may inhibit BMP signaling that is required for fate specification of neural crest-derived pigment cell lineages as previously demonstrated [47]. Another finding in the present study was enhanced mortality induced by exogenous treatment of nApoE4_1-151_, which was not observed in parallel experiments with nApoE3_1-151_. These results support our previous *in vitro* findings indicating enhanced cellular toxicity of only nApoE4_1-151_, which differs by a single amino acid at position 112 (R>C) [17]. The further identification of nApoE4_1-151,_ within the nucleus of neurons of the developing nervous system of zebrafish supports the hypothesis that the uptake of this fragment and trafficking to the nucleus may lead to stimulation of cell death pathways similar to what we have recently observed in BV2 microglia cells [15, 21].

In the context of AD, an important finding of our results was the evolution of tau pathology following exogenous treatment of zebrafish with nApoE4_1-151_ at both 48 and 72 hpf. These findings support that nApoE4_1-151_ may promote tau pathology, a hallmark feature seen in human AD pathology [48]. Previous studies have linked the presence of amino-terminal fragments of ApoE4 with NFTs in the human AD brain as well as in various animal models [14, 25]. The observed tau pathology can lead to disruptions in axonal function which in turn, can have deleterious effects on neuronal signaling and axonal transport [49, 50].

## 5. Conclusion

The *APOE4* allele stands out as the greatest risk factor for late-onset AD, as *APOE4* carriers account for 65-80% of all cases [51]. Although ApoE4 plays a normal role in lipoprotein transport, how it contributes to AD pathogenesis remains speculative. Recent data from our lab suggests that, *in vitro*, nApoE4_1-151_ can traffic to the nucleus leading to toxicity and expression of inflammatory genes in BV2 microglia cells [15, 17, 21]. In the present study, we now expand those findings *in vivo*, by demonstrating toxicity of nApoE4_1-151_, developmental abnormalities, and motor behavior deficits in a novel model system consisting of zebrafish. Taken together, these results support the hypothesis that a key step in mechanism of action is the cleavage of full-length ApoE4, generating an amino-terminal fragment that exhibits a toxic-gain of function. Therefore, the neutralization of this amino-terminal fragment of ApoE4, specifically, may serve as an important therapeutic strategy in the treatment of AD.

## Acknowledgements

This work was funded by the National Institutes of Health Grant 2R15AG042781-02A1. The project described was supported by Institutional Development Awards (IDeA) from the National Institute of General Medical Sciences of the National Institutes of Health under Grants #P20GM103408 and #P20GM109095. We also acknowledge support from the Biomolecular Research Center at Boise State with funding from the National Science Foundation, Grants #0619793 and #0923535; the M.J. Murdock Charitable Trust; and the Idaho State Board of Education.

## Supporting information captions

**Supplemental Figure 1: Tau Pathology present after treatment with exogenous nApoE4**_**1-151**_ **in 48 hpf zebrafish brain. (A-D)**: Representative 40X images from confocal immunofluorescence in 5 μm paraffin embedded sections of non-treated control 48 hpf zebrafish embryos that were stained with DAPI, PHF-1 (1:250), and Anti-6X His Tag antibody to detect nApoE4_1-151_ (1:500). **(E-H):** Parallel experiments as in Panels A-C with the exception that zebrafish embryos were treated exogenously for 24 hours with 25 µg/ml of nApoE3_1-151_. Little to no PHF-1 detected at 48hpf in nApoE3_1-151_. **(I-L):** Parallel experiments as in Panels A-C with the exception that zebrafish embryos were treated exogenously for 24 hours with 25 µg/ml of nApoE4_1-151_. Detection of PHF-1 was much more evident compared to controls and nApoE3_1-151_ sections.

